# In-vivo characterization of magnetic inclusions in the subcortex from non-exponential transverse relaxation decay

**DOI:** 10.1101/2023.09.15.557912

**Authors:** Rita Oliveira, Quentin Raynaud, Ileana Jelescu, Valerij G. Kiselev, Evgeniya Kirilina, Antoine Lutti

## Abstract

According to theoretical studies, the MRI signal decay due to transverse relaxation, in brain tissue with magnetic inclusions (e.g. blood vessels, myelin, iron-rich cells), exhibits a transition from a Gaussian behaviour at short echo times to exponential at long echo times. Combined, the Gaussian and exponential decay parameters carry information about the inclusions (e.g., size, volume fraction) and provide a unique insight into brain tissue microstructure. However, gradient echo decays obtained experimentally typically only capture the long-time exponential behaviour. Here, we provide experimental evidence of non-exponential transverse relaxation signal decay at short times in human subcortical grey matter, from MRI data acquired in vivo at 3T. The detection of the non-exponential behaviour of the signal decay allows the subsequent characterization of the magnetic inclusions in the subcortex.

The gradient-echo data was collected with short inter-echo spacings, a minimal echo time of 1.25 ms and novel acquisition strategies tailored to mitigate the effect of motion and cardiac pulsation. The data was fitted using both a standard exponential model and non-exponential theoretical models describing the impact of magnetic inclusions on the MRI signal. The non-exponential models provided superior fits to the data, indicative of a better representation of the observed signal. The strongest deviations from exponential behaviour were detected in the substantia nigra and globus pallidus. Numerical simulations of the signal decay, conducted from histological maps of iron concentration in the substantia nigra, closely replicated the experimental data – highlighting that non-heme iron can be at the source of the non-exponential signal decay.

To investigate the potential of the non-exponential signal decay as a tool to characterize brain microstructure, we attempted to estimate the properties of the inclusions at the source of this decay behaviour using two available analytical models of transverse relaxation. Under the assumption of the static dephasing regime, the magnetic susceptibility and volume fractions of the inclusions was estimated to range from 1.8 to 4 ppm and from 0.02 to 0.04 respectively. Alternatively, under the assumption of the diffusion narrowing regime, the typical inclusion size was estimated to be ∼2.4 *μ*m. Both simulations and experimental data point towards an intermediate regime with a non-negligible effect of water diffusion to signal decay. Non-exponential transverse relaxation decay provides new means to characterize the spatial distribution of magnetic material within subcortical grey matter tissue with increased specificity, with potential applications to Parkinson’s disease and other pathologies.

## 1 Introduction

The decay of gradient-echo Magnetic Resonance Imaging (MRI) data due to transverse relaxation is widely considered to follow an exponential behaviour with a rate 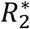 (Weiskopf et al., 2014). Estimates of 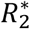 correlate with iron concentration within brain tissue (Péran et al., 2009; Yao et al., 2009; Fukunaga et al., 2010; Langkammer et al., 2010), a property of primary importance for the study of the brain. Iron plays a crucial role in various biological processes such as myelin synthesis, energy production, neurotransmitter synthesis and signalling (Hare et al., 2013; Möller et al., 2019). Therefore some cell types including oligodendrocytes, microglia and dopaminergic neurons exhibit elevated cellular iron concentrations. Abnormal accumulation of iron constitutes a hallmark of neurodegenerative disorders such as Parkinson’s Disease (Gerlach et al., 1994; Thompson et al., 2001; Ward et al., 2014) and can be monitored non-invasively in patients using 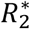 mapping data (Ulla et al., 2013; Damulina et al., 2020). Empirical models have been proposed to link 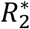 to the overall iron concentration (Schweser et al., 2011; Stüber et al., 2014). However, these models lack a biophysical foundation crucial for specificity, and fail to capture the characteristics of iron-rich cells in the tissue.

Microscopic inclusions of magnetic material within brain tissue such as iron-rich cells, myelin, or blood vessels, induce microscopic inhomogeneities of the magnetic field which can result in a non-exponential gradient-echo signal decay (Haacke et al., 2005; Kiselev and Novikov, 2018). According to theoretical studies, signal decays that result from these magnetic field inhomogeneities display a Gaussian behaviour at short echo times followed by an exponential behaviour at longer echo times (Yablonskiy and Haacke, 1994; Jensen and Chandra, 2000a; Kiselev and Novikov, 2002, 2018; Sukstanskii and Yablonskiy, 2003). Combined, the coefficients that describe the Gaussian and exponential behaviours carry complementary information about those inclusions (e.g., volume fraction, magnetic susceptibility, size). If measured experimentally, these coefficients allow the assessment of the inclusions with improved specificity, offering valuable insights into the cellular underpinnings of neurodegenerative diseases. However, while this non-exponential behaviour has been observed in suspensions of paramagnetic beads (Storey et al., 2015), in ex vivo brain samples (Jensen and Chandra, 2000a), and in vivo blood vessels (Ulrich and Yablonskiy, 2016), no such evidence exists in iron-rich subcortical grey matter.

The characterization of the magnetic inclusions at the source of non-exponential signal decay requires biophysical models of transverse relaxation that establish a quantitative link between the measured MRI data and the properties of the inclusions within brain tissue (e.g. volume fraction, magnetic susceptibility…). These models capture two phenomena. One is the distribution of Larmor frequencies experienced by water molecules due to the microscopic spatial magnetic field inhomogeneities induced by the inclusions (Yablonskiy and Haacke, 1994; Jensen and Chandra, 2000b). These inhomogeneities are static and their effect on the MRI signal is in principle re-focusable using spin-echoes. The other is the temporal effects of water diffusion across this inhomogeneous field - which are non-refocusable (Anderson and Weiss, 1953; Jensen and Chandra, 2000a). Both spatial and temporal effects contribute to signal decay (see Kiselev and Novikov, 2018 for a review). However, analytical expressions of the MRI signal may only be derived from these biophysical models under two mutually exclusive limiting cases. In one case (static dephasing regime, SDR), the spatial inhomogeneities constitute the dominant mechanism underlying signal decay (Yablonskiy and Haacke, 1994; Jensen and Chandra, 2000b). In the other (diffusion narrowing regime, DNR), the temporal effects dominate (Anderson and Weiss, 1953; Kennan et al., 1994; Jensen and Chandra, 2000a; Kiselev and Novikov, 2002; Sukstanskii and Yablonskiy, 2003).

The question of which regime is more suitable to describe the biophysics of transverse relaxation within brain tissue is a topic of debate and depends on the magnetic field strength and brain region under consideration (Sedlacik et al., 2014; Brammerloh et al., 2021; Yablonskiy et al., 2021). In the subcortex, quantitative assessment of iron distribution at the microscopic scale revealed a complex distribution of paramagnetic iron, characterized by a substantial amount dispersed diffusely throughout the tissue, in addition to localized iron-rich cells (Kirilina et al., 2020; Friedrich et al., 2021; Brammerloh et al., 2024). Biophysical models informed by such detailed quantitative measurements have demonstrated that the SDR is suitable to describe the contribution of dopaminergic neurons in the substantia nigra (SN) at field strengths of 7T or above (Sedlacik et al., 2014; Brammerloh et al., 2021). At lower field strengths, diffusion needs to be taken into account (Brammerloh et al., 2024). Because such detailed findings have not been presented for other subcortical nuclei and populations of iron-rich cells, our understanding of which relaxation regime dominates remains fragmented.

In this work, we provide experimental evidence of non-exponential MRI signal decay due to transverse relaxation in subcortical brain regions, in gradient-echo data acquired in vivo at 3T. We fitted the signal decay with an empirical expression and with theoretical models of the effect of magnetic inclusions on the MRI signal (Anderson and Weiss, 1953; Yablonskiy and Haacke, 1994; Jensen and Chandra, 2000a; Sukstanskii and Yablonskiy, 2003). From the value of the Gaussian and exponential parameters of the signal decay, we estimated the properties of the magnetic inclusions under the assumption of the SDR and DNR. For the SN, these estimates were compared with the microscopic distribution of iron known from analyses of ex vivo brain tissue.

## 2 Theory

### 2.1 Transverse relaxation in the presence of magnetic inclusions

Differences between existing biophysical models of the effect of magnetic inclusions on transverse relaxation rely mainly involve the dominating dephasing regime (SDR or DNR) and secondary assumptions about the spatial distribution of magnetic material at the microscopic scale. In particular, models derived in the DNR for weak magnetic field inhomogeneities differ in the form of the auto-correlation function of the Larmor frequency experienced by diffusing spins over time. In the model of Anderson and Weiss (Anderson and Weiss, 1953) (AW), this auto-correlation function is assumed to take an exponential form. In the model of Sukstanskii and Yablonskiy (Sukstanskii and Yablonskiy, 2003) and Jensen and Chandra (Jensen and Chandra, 2000a) (JC), the auto-correlation function was derived analytically from Gaussian water diffusion within the tissue. All existing models predict asymptotic behaviours of the signal decay of the following forms:

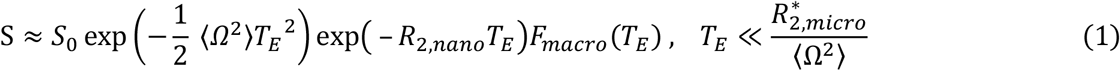

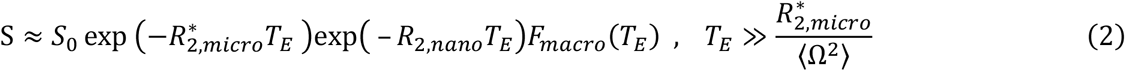

where *S*_0_ is the signal amplitude at *T*_*E*_ = 0, *R*_2,*nano*_ is the transverse relaxation rate due to spin interactions at the molecular/nanoscopic scale, *F*_*macro*_(*T*_*E*_) is the effect of macroscopic magnetic field inhomogeneities (Yablonskiy et al., 2013), ⟨Ω^2^⟩ is the variance of the field inhomogeneities induced by the magnetic inclusions (Novikov et al., 2018) and 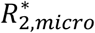 is the transverse relaxation rate induced by the magnetic inclusions at the microscale. A parametric evaluation of these expressions was conducted with a Padé approximation, which is a flexible model-free signal representation (Novikov et al., 2018) derived from a fraction of two polynomials with coefficients adjusted to satisfy the required asymptotic forms at the short- (Gaussian) and long-(exponential) time limits:

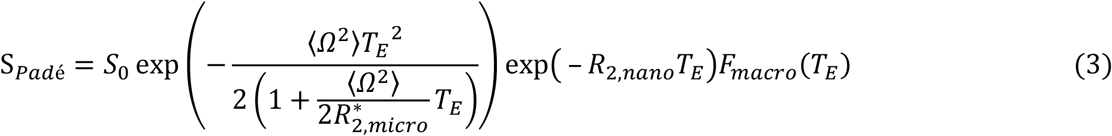

We also used the AW and JC models after parameterization in terms of ⟨Ω^2^⟩ and 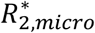:

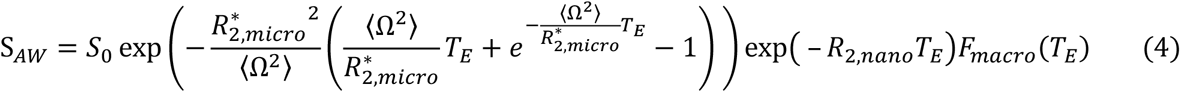

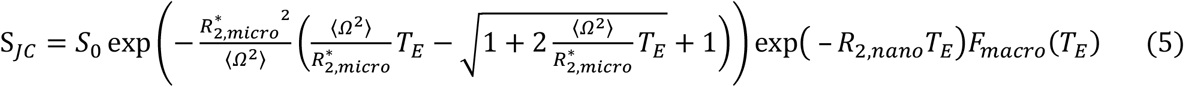

The exponential approximation of the signal is simply:

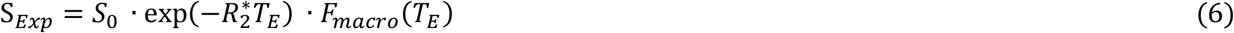

with 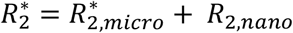.

### 2.2 Microscopic underpinnings of non-exponential decay

The MRI parameters 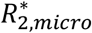 and ⟨Ω^2^⟩ of the signal decay can be linked to the microscopic properties of the inclusions that contain the magnetic material (e.g. iron-rich cells), assumed to have a spherical shape. In particular, the mean square frequency deviation ⟨Ω^2^⟩ of the magnetic field inhomogeneities generated by randomly distributed inclusions is (Yablonskiy and Haacke, 1994; Jensen and Chandra, 2000a; Sukstanskii and Yablonskiy, 2003):

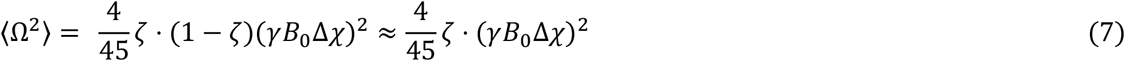

where *ζ* ≪ 1 is the volume fraction of the magnetic inclusions, Δ*χ* is their susceptibility difference with the surrounding tissue (SI units), *γ* the gyromagnetic ratio (2.675 · 10^8^ *rad s*^−1^*T*^−1^) and *B*_0_ the main magnetic field. Note that ⟨Ω^2^⟩ is a measure of magnetic field inhomogeneities averaged across an imaging voxel. By contrast, the characteristic Larmor frequency induced by a single sphere is: 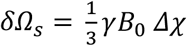 (Yablonskiy and Haacke, 1994).

In the framework of the SDR (Yablonskiy and Haacke, 1994) and DNR (Jensen and Chandra, 2000a; Sukstanskii and Yablonskiy, 2003), the relaxation rate is described by the following equations:

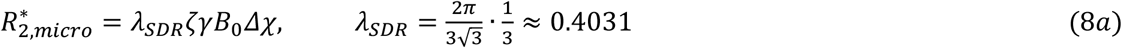

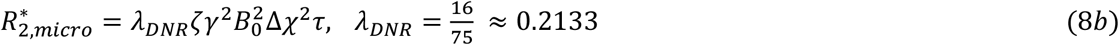

where 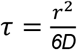 is the time scale for water molecules to diffuse away from a spherical magnetic inclusion of radius *r. D* is the water diffusion coefficient in tissue (1 μm^2^/ms).

In the DNR, the dimensionless parameter *α* = *τ* · *δΩ*_*s*_ ∝ *r*^2^ (Yablonskiy and Haacke, 1994; Kiselev and Novikov, 2002) represents the amount of spin dephasing induced by the field inhomogeneities over the period *τ*. In the DNR, the condition *α*≪1 must be verified. The DNR may therefore apply to distributions of magnetic inclusions with comparatively smaller sizes than the SDR. Note that the relaxation rate of the DNR is parametrically smaller than that of the SDR: 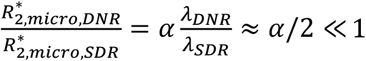. As a result, the relaxation rate of the MRI data will yield very different properties of the inclusions (*Δχ, ζ τ*) under the SDR and DNR.

## 3 Methods

### 3.1 Participant cohort

MRI data were acquired from 5 healthy volunteers (2 females, mean age=32±9 years old). The study received approval by the local ethics committee and all volunteers gave written informed consent for their participation.

### 3.2 Data acquisition

MRI data were acquired on a 3T whole-body MRI system (Magnetom Prisma; Siemens Medical Systems, Erlangen, Germany) with a 64-channel receive head coil and a custom-made multi-echo 3D fast low-angle shot (FLASH) pulse sequence with bipolar readout. To facilitate the detection of a non-exponential signal decay, 16 gradient-echo images were acquired with a minimal echo time of 1.25 ms, and inter-echo spacing of 1.2 ms. The radio-frequency (RF) flip angle was 12° and the repetition time was 23.2 ms. Image resolution was 1.2 mm^3^ isotropic, with a field of view 208×192×144 mm along the read and two phase-encode directions. Partial Fourier (factor 6/8) was used in the phase and partition directions. Three repetitions were conducted on each participant and the total nominal acquisition time was 18min09s.

To minimize image degradation due to head motion, an optical tracking prospective motion correction system (KinetiCor, HI, Honolulu) was used (Zaitsev et al., 2006; Maclaren et al., 2012). Cardiac pulsation constitutes an additional source of noise in brain relaxometry data which accounts for up to 35% of the variability of 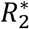 maps across repetitions (Raynaud et al., 2023). To minimize the effect of cardiac-induced noise, the cardiac pulsation of the participants was recorded using a finger pulse oximeter and data acquisition was suspended during the systolic period of the cardiac cycle, taken to last for a duration of 300 ms (Raynaud et al., 2023). For a heart rate of 80 beats per minute, this strategy resulted in an increase in scan time by approximately 40%. As a result of these prospective strategies for the correction of head motion and cardiac pulsation, the motion degradation index (Castella et al., 2018; Lutti et al., 2022; Corbin et al., 2023), an index of data quality, lied within a narrow range across participants and did not exceed 3.4 s^−1^ (Figure S1).

Multi-parameter mapping (Weiskopf et al., 2013) data was acquired to compute maps of the MRI parameter MTsat (magnetization transfer saturation), a semi-quantitative parameter reflecting tissue myelination with improved contrast between tissue classes, allowing an accurate delineation of subcortical grey matter regions (Helms et al., 2009). The protocol comprised three multi-echo 3D FLASH scans acquired with magnetization transfer-, proton density- and T1-weighting (RF excitation flip angle = 6º, 6º and 21º, respectively; repetition time TR=24.5 ms). Eight echo images were acquired for the T1- and proton density-weighted contrasts and six for the magnetization transfer-weighted contrast. Image resolution was 1 mm^3^ isotropic, and the image field of view was 176×240×256 mm. B1-field mapping data was acquired (4 mm^3^ voxel size, TR/TE = 500/39.1 ms) to correct RF transmit field inhomogeneity effects on the MTsat maps (Lutti et al., 2010, 2012). For correction of image distortions in the B1 map data, B0-field map data was acquired with a 2D double-echo FLASH, TR=1020 ms, α =90°, TE1/TE2 = 10/12.46 ms, BW = 260 Hz/pixel, slice thickness = 2 mm. The motion correction system described above was also used here.

### 3.3 Anatomical imaging processing

MTsat maps were calculated from the magnetization transfer-, proton density- and T1-weighted images with the hMRI toolbox (https://hMRI.info) (Tabelow et al., 2019), as described in (Helms et al., 2008a, 2008b; Weiskopf et al., 2013). MTsat maps were segmented into grey and white matter tissue probability maps using the Statistical Parametric Mapping software (SPM12, Wellcome Centre for Human Neuroimaging, London) (Ashburner and Friston, 2005). A grey matter mask was computed by selecting voxels with a grey matter probability above 0.95. Globus pallidus (GP), putamen, thalamus, and caudate regions of interest (ROI) were defined from the grey matter mask and the regional labels of the Neuromorphometrics atlas (http://neuromorphometrics.com/). As no label exists for the SN, this region was delineated using an *ad hoc* procedure, from a cuboid placed appropriately in the space of each MTsat map. Within this cuboid, SN voxels were identified from the grey matter voxels labelled as brainstem and ventral diencephalon in the Neuromorphometrics atlas. Beyond subcortical grey matter, the fusiform gyrus was also defined from the grey matter mask and the regional label of the Neuromorphometrics atlas, serving as a reference region with a low concentration in non-heme iron (Haacke et al., 2005).

### 3.4 Fitting of the transverse relaxation decay

Data were analyzed using bespoke analysis scripts written with MATLAB R2021a (The Mathworks, Natick, MA). The effect of macroscopic magnetic field inhomogeneities on the gradient-echo signal (*F*_*macro*_ in Eqs. 1-6) was corrected with the voxel spread function (VSF) method (Yablonskiy et al., 2013). The complex gradient-echo data were denoised using the Marchenko Pastur-PCA method (Veraart et al., 2016b, 2016a; Does et al., 2019), using cubic regions of 5×5×5 voxels. At each voxel, we removed scaling and additive effects between the signal decays acquired across repetitions, due to e.g. head motion in the spatially varying sensitivity profile of the receive coil. To suppress the noise floor in the magnitude images, background voxels outside the head were identified from the segmentation of the first gradient-echo image using SPM12 (Ashburner and Friston, 2005). The distribution of signal intensities across noise voxels was fitted assuming a Rician distribution and the resulting value of the noncentrality parameter was deducted from the signal intensities.

Fitting of the transverse relaxation decay with the analytical expressions of Section 2.1 was conducted using non-linear least square minimization (*lsqnonlin* Matlab function) with a trust-region-reflective algorithm. 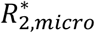 was bounded between 1 to 80 s^-1^ with an initial value of 20 s^-1^ for the signal models of Eqs 3-5. The ⟨Ω^2^⟩ parameter ranged from 100 to 4×10^4^ *rad*^2^*s*^−2^ for the Padé and AW models and from 100 to 8×10^4^ *rad*^2^*s*^−2^ for the JC model with an initial value of 10^4^ *rad*^2^*s*^−2^ for all of them. 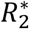 from Eq. 6 ranged from 0 to 80 s^-1^ with an initial value of 20 s^-1^. *S*_0_ was bounded between 10 and 2000 with an initial value of 500. *R*_2,*nano*_ was not estimated during data fitting due to the unsuitable range of echo times of the data and was set to 10 s^-1^ instead (Jensen and Chandra, 2000a; Sedlacik et al., 2014; Brammerloh et al., 2021). As *R*_2,*nano*_ depends on tissue iron concentration, this carries the risk of misattributing the actual value of *R*_2,*nano*_ to 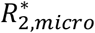.

While the Padé expression is not strictly speaking a model but a representation of the MRI signal, we henceforth refer to all three analytical expressions (Eqs. 3-5) as models of the MRI signal for the sake of simplicity. The goodness of fit of each model was estimated from the mean squared error of the fit (MSE) and the Akaike Information Criterion (AIC), which includes a penalty for model complexity. Lower MSE and AIC values indicate a better model fit. Model parameter estimates for the five regions of interest were extracted from all voxels and all subjects after the removal of the voxels with high MSE (>15), indicative of spurious effects in the data such as physiological noise (as reference the average MSE across all voxels is ∼8). We also excluded voxels where the transition from Gaussian to exponential behaviour took place over a timescale 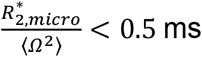 after RF excitation, too short to be robustly detectable.

### 3.5 Microscopic underpinnings of non-exponential decay

From the estimates of 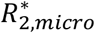 and ⟨Ω^2^⟩ et each voxel, we attempted to estimate the properties of the magnetic inclusions at the source of the non-exponential decay within brain tissue. This analysis was conducted under the two mutually exclusive scenarios of the SDR and DNR, each providing a different interpretation of the decay curve parameters in terms of microstructural tissue properties.

Under the assumption of the SDR, we estimated the magnetic susceptibility (Δ*χ*) and volume fractions (*ζ*) of the inclusions (Eqs. 7 and 8a). Under the assumption of the DNR, where diffusion effects are considered, the model parameters 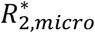 and ⟨Ω^2^⟩ depend not only on Δ*χ* and *ζ* as in the SDR, but also on *τ*. Since all three properties cannot be estimated separately from 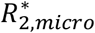 and ⟨Ω^2^⟩ alone, only *τ* was estimated (Eqs. 7 and 8b).

We conducted non-parametric Kruskal-Wallis statistical tests of inter-regional differences in the estimates of Δ*χ* and *ζ* obtained under the assumption of the SDR (*kruskalwallis* function in Matlab 2021). Post-hoc Tukey’s HSD tests were conducted subsequently to identify the pairs of regions at the source of these differences (*multcompare* function in Matlab). The effect size was computed using the cliff’s delta to quantify the magnitude of differences between regions.

### 3.6 Non-heme iron as a possible source of the non-exponential signal decay

To investigate if the detected non-exponential decay can be induced by microscopic inclusions of non-heme iron with cellular sizes, we numerically simulated the gradient-echo signal decay induced by iron-rich cells of the SN at 3T. The numerical simulations were conducted from the cellular distribution of iron in neuromelanin-pigmented dopaminergic neurons within the volume of a typical MRI voxel, quantified in 3D with microscopic resolution from a post-mortem brain (Brammerloh et al., 2021), and estimates of the magnetic susceptibility of iron and neuromelanin determined using a combination of MRI microscopy and micro X-ray fluorescence (Brammerloh et al., 2024).

The 3D iron distribution was converted into magnetic susceptibility maps using the estimates of neuromelanin magnetic susceptibility obtained experimentally and literature values for ferritin, as described in (Brammerloh et al., 2021, 2024). These maps were convolved with a magnetic dipole kernel to compute the Larmor frequency distribution within this voxel due to the presence of iron and neuromelanin. We then used two methods to simulate the gradient-echo signal decay generated by this frequency distribution: 1) the SDR approximation; and 2) Monte Carlo simulations accounting for water diffusion using a typical water diffusion coefficient of brain tissue of 1 µm^2^/ms. The decay resulting from the Monte Carlo simulations was fitted with an exponential for TE > 10 ms and with Eq. 3 for all echoes.

## 4 Results

### 4.1 Non-exponential transverse decay in subcortical tissue

At short echo times (*T*_*E*_ <5-10ms), transverse signal decays in the basal ganglia and thalamus (Figure 1) display systematic deviations from exponential behaviour: the logarithm of the signal exhibits an initial quadratic form, with a transition towards a linear dependence only at longer times. This behaviour is consistent with the effect of magnetic inclusions (e.g iron-rich cells, myelin, or blood vessels) on the MRI signal predicted by theoretical studies (see (Kiselev and Novikov, 2018) for a review).

**Figure 1.**
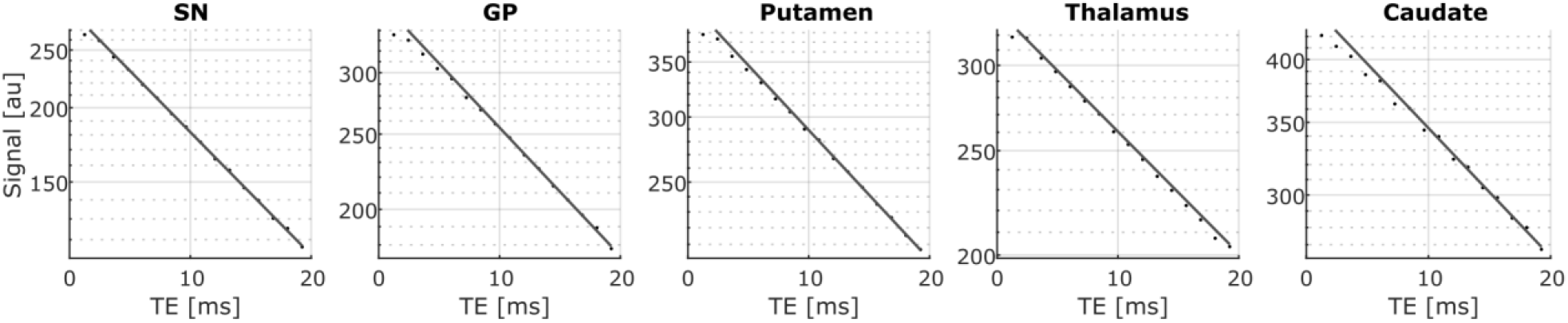
Transverse relaxation decays in one representative voxel of each subcortical grey matter region (semilog-scale). The line shows the exponential decay fit at long echo times (T_E_>10ms). At short echo times (TE≲5ms), the data deviates from the exponential decay fit (line), displaying a quadratic decay consistent with the effect of magnetic inclusions on the MRI signal predicted by the theory.

The non-exponential models of transverse relaxation (Padé, AW, JC) can account for the non-exponential decay of the MRI signal at short times, leading to an improved fit of the data (Figure 2A). The residual levels are largely consistent across the non-exponential models (MSE ∼5), a factor of ∼1.6 smaller than for the exponential fit (MSE ∼8) (Figure 2B). Similarly, the AIC decreases by a factor of ∼1.2 between the exponential (AIC ∼100) and non-exponential fits (AIC∼90) (Figure 2C), showing that the residual decrease goes beyond that expected from the higher number of parameters of non-exponential models.

**Figure 2.**
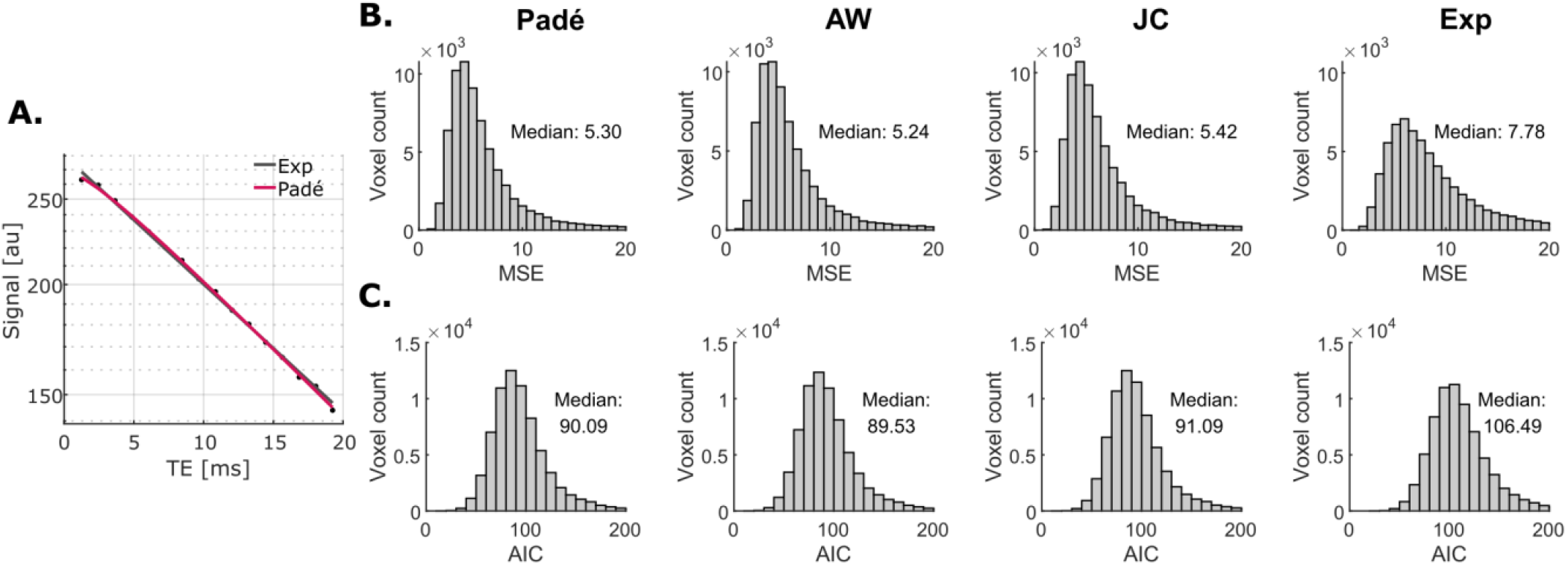
Residual levels across signal models for all subcortical regions analyzed. Example fit for a representative voxel in the SN (semilog-scale). Non-exponential models of transverse relaxation (Padé, AW, JC) can account for the non-exponential decay at short times, leading to an improved fit of the data (A). As a result, the median of residual levels (MSE) is reduced by ∼1.6 across subcortical regions, consistently for all non-exponential models (B). This residual decrease leads to a decrease of the median AIC by ∼1.2, beyond that expected from the higher number of parameters of non-exponential models (C).

The ratio of the MSE between the exponential and non-exponential fits is higher in subcortical regions (average ∼1.6) than in cortical grey matter (average ∼1.4, Figure 3A and 3B), showing that the non-exponential behaviour takes place predominantly there. In particular, this ratio takes a value of 2 in the iron-rich GP and a value of 1.3 in the iron-poor fusiform gyrus (Haacke et al., 2005). Figure 3C shows example signal decays from these two regions.

**Figure 3.**
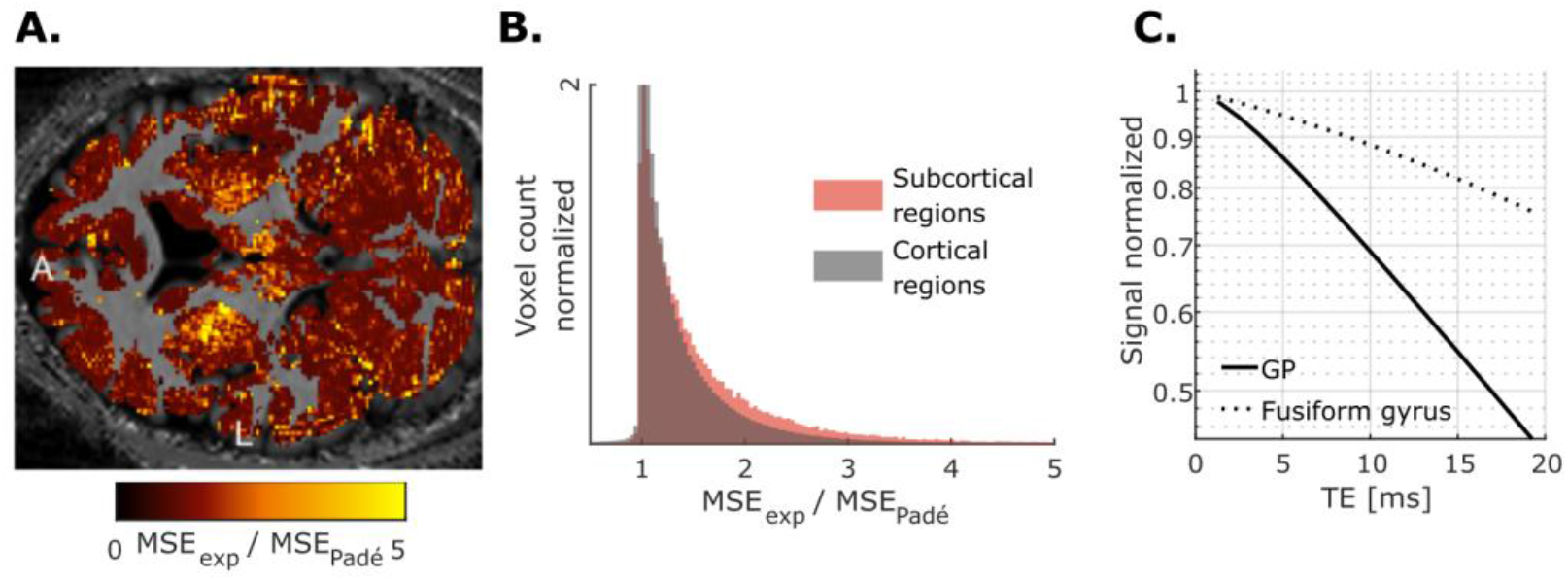
Spatial distribution of the ratio of the MSE obtained from the fits with the exponential and Padé signal models (A). The higher ratio values in subcortical regions (average ∼1.6) indicate that stronger deviations from exponential behaviour take place in these areas (B). The stronger non-exponential behaviour in the iron-rich GP than in the iron-poor fusiform gyrus is illustrated for a representative voxel (C, semilog-scale). L – left; A – anterior.

### 4.2 Estimates of the MRI signal model parameters

Non-exponential signal decays were reliably detected with 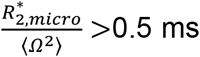 in 71/81/82/83/77/18% of voxels in the SN/GP/putamen/thalamus/caudate/fusiform gyrus, respectively. In these voxels, the non-exponential models (Padé, AW, JC) lead to estimates of 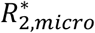 that are respectively 37%, 30%, and 54% higher than the exponential approximation (Figure 4A).

**Figure 4.**
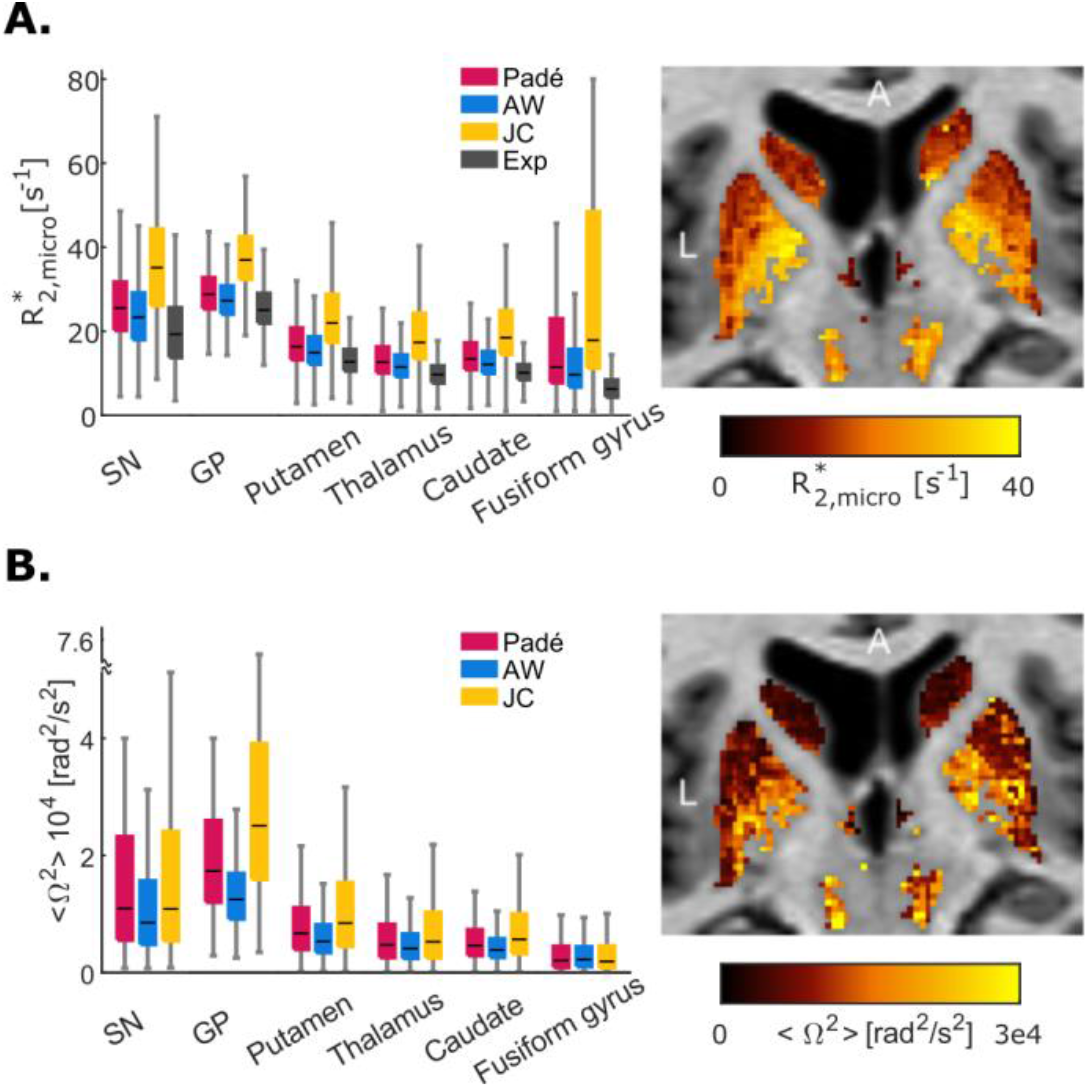
Non-exponential model parameter estimates. Estimates of 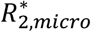 (A) and ⟨Ω^2^⟩ (B) are highest in the GP followed by the SN and lowest in the fusiform gyrus, in agreement with histological measures of iron concentration (Hallgren and Sourander, 1958) and Eqs. 7 and 8. The example maps of 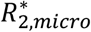 and ⟨Ω^2^⟩ were obtained from the AW model. L – left; A – anterior; Cau – caudate; Put – putamen; GP – globus pallidus; Thal – thalamus; SN – substantia nigra.

The estimates of 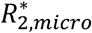 and ⟨Ω^2^⟩ are spatially organized and are consistent between anatomical regions (Figure 4). The values of 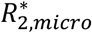 and ⟨Ω^2^⟩ are higher in the GP and the SN, and lowest in the fusiform gyrus, in agreement with histological measures of iron concentration in brain tissue (Hallgren and Sourander, 1958) and with the expected dependence of these parameters on iron content (Eqs. 7 and 8).

The JC model yields systematically higher values of 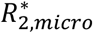 and ⟨Ω^2^⟩ than the AW model and Padé approximation (∼31-73%). This arises from the square-root term in the JC signal equation (Eq. 5), which introduces a broad modulation of the signal over the range of echo times of the data (TE < 20 ms), i.e. a slow transition to a monoexponential decay. As a result, the decay rate of the data at TE ∼10-20ms differs from the estimates of 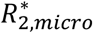 at TE → ∞ provided by the JC model: the decay of the data and that of the exponential part of the JC fit have different slopes (Figure 5). On the other hand, the estimates of 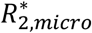 from the AW model match the decay rate of the data at TE ∼10-20ms. Results from the Padé approximation and the AW model are largely consistent.

**Figure 5.**
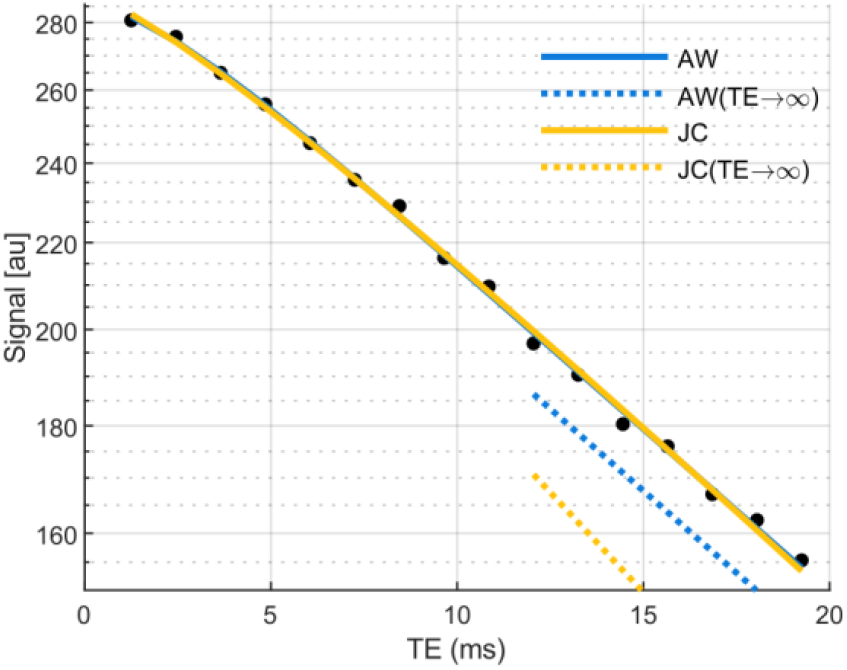
Example transverse relaxation decay from a voxel in the SN and corresponding fits with the AW (blue solid line) and JC (yellow solid line) models (semi-log scale). For each model, the asymptotic behaviour (TE → ∞) plotted as in (Kiselev and Novikov, 2018) (dashed lines), reflects the estimates of 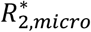. For the AW model, the slope of the experimental data at TE ∼10-20ms matches that of the asymptotic behaviour. This is not the case for the JC model due to the square-root term in the JC signal equation: the 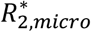 estimates provided by the JC model are higher than the decay rate of the data at TE ∼10-20ms.

### 4.3 Non-heme iron as a possible source of the non-exponential signal decay

Post-mortem studies of the microscopic cellular distribution of iron in SN demonstrate that dopaminergic neurons accumulate high levels of iron stored in neuromelanin (Figure 6A) (Brammerloh et al., 2021). In adult human brains, these neuromelanin clusters are about 15 *μ*m in radius and contain approximately 300-1000 *μ*g/g of iron (Figure 6A). At 3T, the Larmor frequency perturbations that arise from these paramagnetic inclusions (Figure 6B) lead to a frequency distribution with a width of ΔΩ ∼35 *rad s*^−1^ (Eq. 7) across the volume of an MRI voxel (Figure 6C). The gradient-echo signal decay that results from these field inhomogeneities using Monte Carlo simulations or the SDR, exhibits a deviation from an exponential behaviour at short echo times (TE < 5 ms, see Figure 6D) consistent with the experimental data (Figure 1). At long echo times, the Monte Carlo simulated signal decay differs from the SDR predictions, suggesting that diffusion effects cannot be ignored for dopaminergic neurons in the SN at 3T. The Monte Carlo simulated decay is better fitted with the Padé model 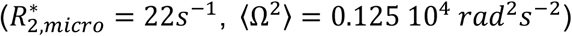 than with an exponential, similar to our experimental data (Figure 2,4).

**Figure 6.**
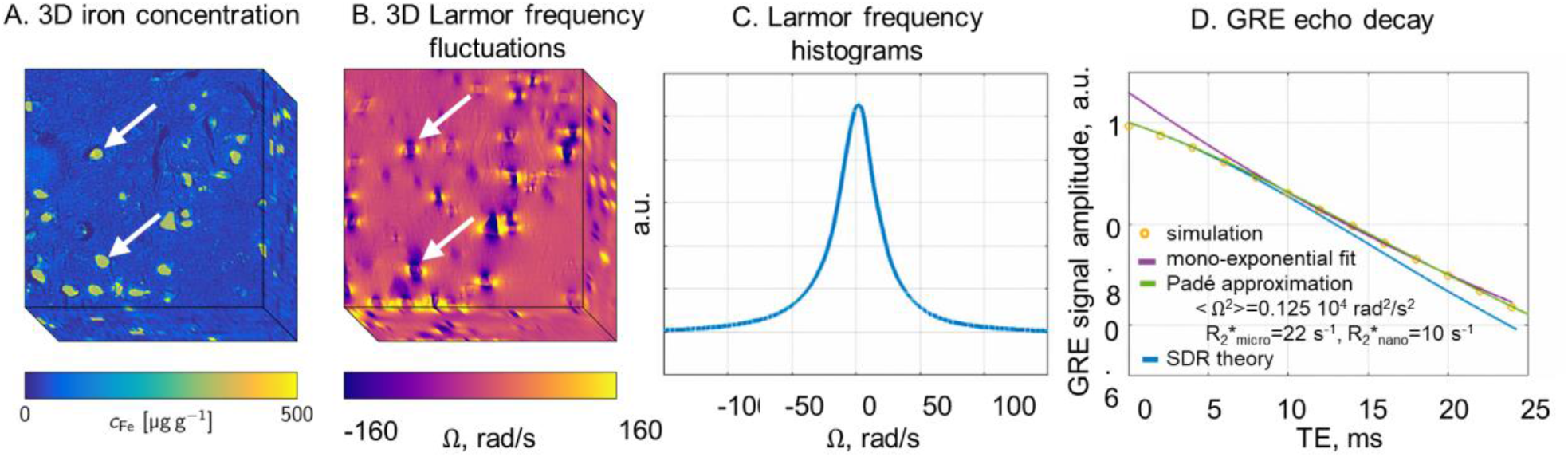
Iron-rich dopaminergic neurons lead to non-exponential signal decay in the SN at 3T. Microscopic iron concentration map obtained from a post-mortem brain (A, adapted from (Brammerloh et al., 2021) under the [Creative Commons Attribution 4.0 International License](https://creativecommons.org/licenses/by/4.0/legalcode.en)), featuring hotspots of iron accumulation inside dopaminergic neurons, as well as diffusely distributed iron outside the neurons. These paramagnetic inclusions lead to local inhomogeneities of the Larmor frequency (B, adapted from (Brammerloh et al., 2021) under the [Creative Commons Attribution 4.0 International License](https://creativecommons.org/licenses/by/4.0/legalcode.en)). As a result, a distribution of ΔΩ ∼35 *rad s*^−1^ of the Larmor frequency takes place across the volume of an MRI voxel (C). The resulting Monte Carlo (circles) and SDR (blue line) simulated gradient-echo signal decays (D) deviate from an exponential at short echo times (TE<5ms), consistently with the experimental data (Figure 1). At long echo times (TE>10 ms), neglecting water diffusion (SDR) leads to a higher exponential decay rate than when water diffusion is accounted for (Monte Carlo simulation). The Padé approximation (green line) provides a better fit to the Monte Carlo simulated signal 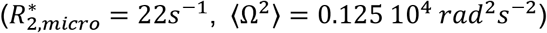 than the exponential model (purple line).

### 4.4 Characterization of magnetic inclusions within subcortical tissue

From the estimates of the MRI signal parameters 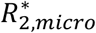 and ⟨Ω^2^⟩, we characterized the properties of the magnetic inclusions present within brain tissue, at the source of the non-exponential decay under two mutually exclusive scenarios – the SDR and DNR.

Scenario 1: Under the assumption of the SDR (Figure 7), the estimates of 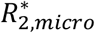 and ⟨Ω^2^⟩ were used to estimate the volume fraction (*ζ*) and magnetic susceptibility (Δ*χ*) of the inclusions (Eqs. 7 and 8a). In subcortical grey matter, the median value of Δ*χ* ranges from 1.8 to 4.0 ppm - largest in the GP and SN (∼2.6 ppm and 2.4 ppm respectively for the AW signal model), followed by the putamen and thalamus (Δ*χ*∼2.0 ppm) and caudate (Δ*χ*∼1.8 ppm). The fusiform gyrus of the cerebral cortex yields the lowest values of Δ*χ* (∼1.5 ppm). The GP and SN show the largest values of *ζ* (median: 0.034 and 0.030 from the AW signal model respectively), followed by the putamen (0.023), caudate (0.021), fusiform gyrus (0.018) and thalamus (0.016). The JC model yields estimates of Δ*χ* that are similar to those of the Padé model and approximately 25% higher than those from the AW model. Additionally, the JC model yields estimates of *ζ* that are about 54% higher compared to the Padé model and approximately 85% higher compared to the AW model. These differences are due to the systematic differences in the 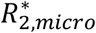 and ⟨Ω^2^⟩ estimates highlighted above.

**Figure 7.**
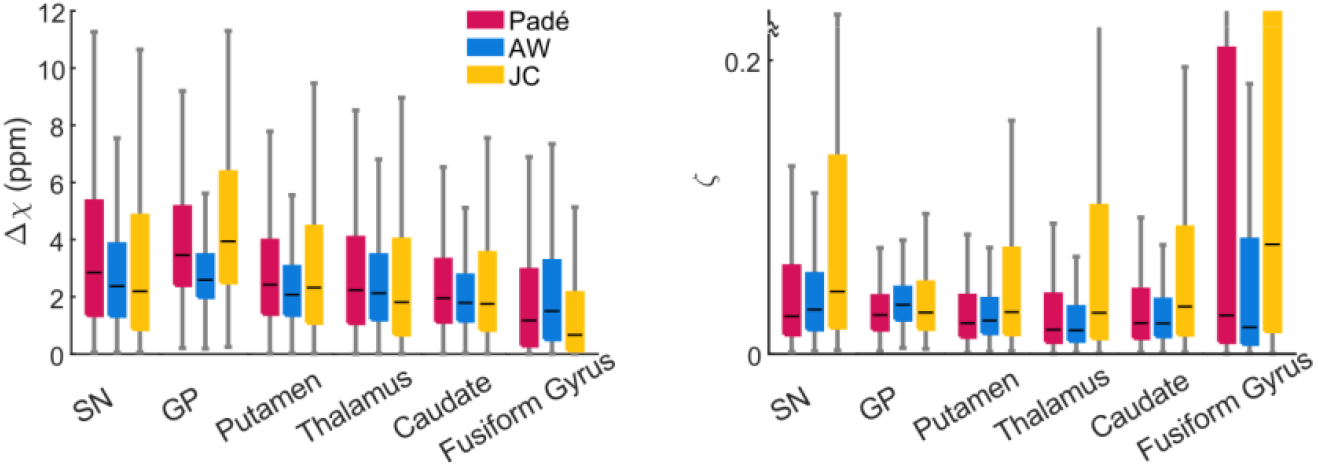
Magnetic susceptibility and volume fractions of magnetic inclusions within brain tissue, estimated from the values of 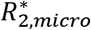 and ⟨Ω^2^⟩ under the assumption of the SDR. Magnetic susceptibilities are larger in the SN and GP (median Δ*χ*∼2.5 ppm) compared to the remaining regions (Δ*χ*∼1.6 ppm). The volume fraction of the inclusions (*ζ*) ranges between ∼0.016 and 0.043 across subcortical regions.

The Kruskal-Wallis tests revealed statistically significant differences in Δ*χ* between at least two ROIs (F(5,76596) = [1579], p < 0.001 for the AW model). The corresponding Tukey’s HSD test for multiple comparisons found significant differences between Δ*χ* estimates from all ROIs (p<0.01) with small effect sizes, except between the thalamus and putamen where no significance was found. The fusiform gyrus showed the strongest effect sizes (0.12-0.35) compared to the remaining regions.

Similarly, the Kruskal-Wallis tests revealed statistically significant differences in *ζ* between at least two ROIs (F(5,76596) = [1744], p < 0.001 for the AW model). The corresponding Tukey’s HSD test showed significant differences between the *ζ* estimates from the GP and those from the putamen, thalamus, and caudate and between SN and thalamus with the largest effect size (>0.30). Other inter-regional differences were found significant (p<0.01) with small effect sizes (<0.20). A detailed overview of this statistical evaluation can be found in Figure S2.

Scenario 2: Under the assumption of the DNR, which for a given 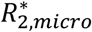 and ⟨Ω^2^⟩, requires a greater total amount of iron compared to the SDR, we estimated the parameter *τ* according to Eqs. 7 and 8b (Figure 8). The estimates of *τ* are larger in the fusiform gyrus (median ∼2.0 ms, from the Padé signal model), followed by the putamen, thalamus, and caudate (median ∼1.0 ms), and finally the SN and GP (median ∼0.8 ms). A value of *τ*∼1.0 ms implies a value of ∼2.4 *μ*m for the magnetic field inhomogeneities generated by the inclusions. In the fusiform gyrus, this radius is ∼3.5 *μ*m (*τ*∼2.0 ms). The JC model yields estimates of *τ* higher than the Padé model by 26% and than the AW model by 35%, due to the systematic differences in the 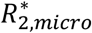 and ⟨Ω^2^⟩ estimates highlighted above.

**Figure 8.**
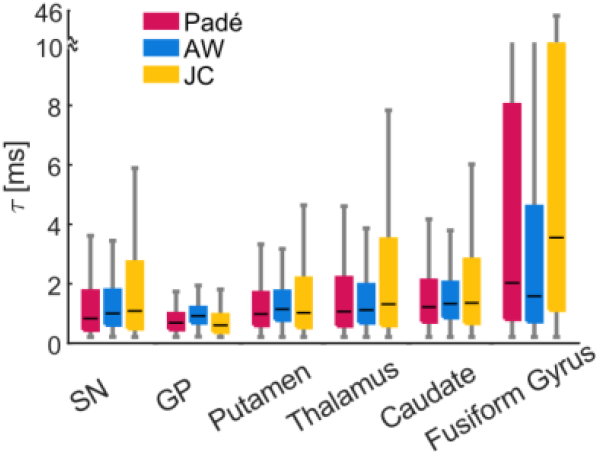
Estimates of the parameter *τ*, computed from the values of 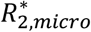 and ⟨Ω^2^⟩ under the assumption of the DNR. The characteristic diffusion times *τ* are larger in the putamen, thalamus, and caudate (median ∼1.0 ms) compared to SN and GP (∼0.8 ms). The fusiform shows the largest values of *τ* (∼2.0 ms).

## 5 Discussion

Here we provide experimental evidence of non-exponential signal decay due to transverse relaxation in *in vivo* MRI data from subcortical regions. This signal decay follows a Gaussian behaviour at short echo times with a transition to exponential behaviour at long echo times. This non-exponential decay is in agreement with theoretical models of the effect of microscopic magnetic inclusions, located within brain tissue, on the MRI signal. These models show an improved fit to the data compared to the widely used exponential model. The strongest deviations from exponential behaviour are found in iron-rich areas such as the GP and SN. From the values of the model parameters, we estimated the properties of the magnetic inclusions. Numerical simulations of the gradient-echo signal from post-mortem maps of iron-rich dopaminergic neurons in SN show that cells rich in non-heme iron can be at the source of this decay behaviour. These results illustrate how non-exponential transverse relaxation signal decay can be used to characterize iron-rich microscopic inclusions from in vivo MRI data.

### 5.1 Non-exponential transverse relaxation decay

The lack of evidence for non-exponential signal decay in subcortical regions has been attributed to the short timescale of the transition between the Gaussian and exponential behaviours, below the range of achievable echo times (Yablonskiy et al., 2021). Here, we combined a dense sampling of the decay curve with acquisition strategies that mitigate the level of physiological noise in the data (Castella et al., 2018; Raynaud et al., 2023) to enable the reliable detection of the non-exponential decay curve. Transverse relaxation decay exhibits a Gaussian behaviour at short echo times (T_E_≲5ms) with a transition towards exponential decay (Figure 1), consistent with theoretical models of the effect on the MRI signal of magnetic inclusions within brain tissue (Kiselev and Novikov, 2002; Novikov and Kiselev, 2008). These models show an improved fit to the data compared with the widely used exponential model (Figure 2). This behaviour was predominantly observed in subcortical grey matter regions (Figure 3), known for their elevated non-heme iron content. In particular, the strongest non-exponential behaviours (i.e. higher values of 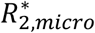 and ⟨Ω^2^⟩) were observed in GP and SN, the areas with the highest iron content (Figure 4) (Hallgren and Sourander, 1958; Haacke et al., 2005; Langkammer et al., 2012; Krebs et al., 2014). While heme-iron in blood vessels may also contribute to the non-exponential decay, information on its spatial distribution within the tissue remains scarce. Myelin, on the other hand, may have a particularly pronounced effect on the thalamus due to its comparatively high myelin content and low iron concentrations (Hallgren and Sourander, 1958).

Transverse relaxation decays obtained from numerical simulations of the gradient-echo signal from post-mortem maps of iron-rich neurons in SN showed the same features as the MRI data acquired experimentally. These findings underscore the contribution of cells rich in non-heme iron to non-exponential signal decay in subcortical grey matter.

We considered a model-free Padé approximation and two models of the MRI signal generated by brain tissue with magnetic inclusions. All corresponding expressions fitted the data equally well, with marginal differences between them (Figure 2). Nonetheless, the JC model yields higher estimates of 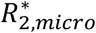 and ⟨Ω^2^⟩ compared to the AW model and Padé approximations. This discrepancy originates from the long transition from the Gaussian to exponential behaviours predicted by the JC model, due to the square-root term in Eq. 5. As a result, the estimates of 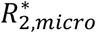 from the JC model differ from the decay rate of the data at TE∼10-20ms (Figure 5).

### 5.2 Characterization of magnetic inclusions within subcortical tissue

From the estimates of the two signal model parameters (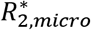and ⟨Ω^2^⟩), we attempted to estimate the properties of the magnetic inclusions embedded within brain tissue, which would not have been achievable from the exponential tail of the decay alone. This was conducted under two mutually exclusive scenarios 1) the SDR (Figure 7) and 2) the DNR (Figure 8), each with distinct analytical expressions to quantitatively link the signal model parameters with the properties of the underlying magnetic inclusions.

Scenario 1: Under the assumption of the SDR, the estimates of the magnetic susceptibility of the inclusions, lie in the range Δ*χ*∼1.8-4 ppm across subcortical regions and models of the MRI signal. The largest values of Δ*χ* are encountered in the GP and SN, followed by the putamen, thalamus and caudate in decreasing order (2.6, 2.4, 2.0, 2.0, 1.8 ppm respectively for the AW model). This ordering follows that of ex vivo measures of bulk iron concentration within the tissue (*C*_*GP*_ ≥ *C*_*SN*_ > *C*_*Putamen*_ > *C*_*Caudate*_ > *C*_*Thalamus*_) (Hallgren and Sourander, 1958; Griffiths et al., 1999; Haacke et al., 2005).

The estimates of Δ*χ* derived from the MRI data can be compared with those obtained from ex vivo studies of non-heme iron distribution. In the SN we can assume that the non-exponential relaxation is induced mainly by iron bound to neuromelanin in dopaminergic neurons. In this case, Δ*χ* = *ρ* · *χ*_*eff_NM*_ · [*Fe*]_*NM*_, where *ρ*∼1.05 g/cm^3^ (Barber et al., 1970) and *χ*_*eff_NM*_∼2.98 ppm m^3^/kg is the effective magnetic susceptibility of neuromelanin (Brammerloh et al., 2024). Taking [*Fe*]_*NM*_∼0.49 mg/g for the concentration of iron in the neuromelanin of dopaminergic neurons (Brammerloh et al., 2021, 2024; Friedrich et al., 2021) one gets: Δ*χ*∼1.5 ppm. These estimates are consistent with the ones obtained from our MRI data (2.4 ppm with the AW model). Since quantitative histological data for other regions are unavailable, we cannot verify the plausibility of our model estimates in those areas. The estimates of Δ*χ* derived from the MRI data differ from the magnetic susceptibility of heme iron in blood capillaries (0.4 to 0.5 ppm (Schenck, 1992; Spees et al., 2001)).

The estimates of the volume fraction of the inclusions computed from the MRI data lie in the range *ζ*∼ 0.02-0.04 across regions and MRI signal models. In particular, in the SN they are ∼0.03 (with the AW model), close to those estimated from histological analyses of the dopaminergic neuron’s volume fraction (0.03 to 0.12 (Brammerloh et al., 2021)). The MRI-derived estimates of *ζ* are also in line with the human capillary vascular volume fraction (∼0.02-0.025 (Buschle et al., 2018)).

Scenario 2: Under the assumption of the DNR, we estimated the parameter *τ*, the decay time of the frequency auto-correlation of water molecules diffusing through the inhomogeneous magnetic field generated by the inclusions. The *τ* estimates (∼1.0 ms) in subcortical grey matter suggest a typical radius *r* of ∼2.4 *μ*m for the magnetic inclusions. This estimate is consistent with the radius of the spherical inclusions reported in other MRI relaxometry studies of excised human grey matter tissue (Jensen and Chandra, 2000a). In particular, the latter study also reported larger values of *r* in the putamen (3.1 *μ*m), thalamus (3.0 *μ*m), and caudate (2.9 *μ*m), compared to the GP (2.3 *μ*m) as observed here (Figure 8). The MRI-derived estimate of *r* is also in the order of a small capillary size (∼3.2 *μ*m (Lauwers et al., 2008)). However, it is also lower than the typical radius of neuronal or glial cells (5-20 *μ*m in neurons, 5-10 *μ*m in microglia, 2.5-10 *μ*m in astrocytes, 2-5 *μ*m oligodendrocytes (Ward et al., 2014; Reinert et al., 2019; Brammerloh et al., 2021; Friedrich et al., 2021)). The separate estimation of Δ*χ, ζ* and *τ* in the DNR, which can involve estimates of MRI susceptibility in addition to the MRI parameters 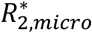 and ⟨Ω^2^⟩ used here, may help clarify the validity of this assumption. Indeed, the condition *α* = *τ* · *δΩ*_*s*_ ≪ 1 must be verified in the DNR. Also, because the relaxation rate of the DNR is parametrically smaller than that of the SDR 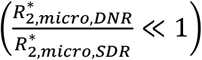, the resulting estimates of Δ*χ* and *ζ* may differ greatly from those presented here.

In conclusion, while the SDR may be plausible in the SN, this may not be the case in other regions. Instead, the most plausible scenario may be that transverse relaxation results from an intermediate dephasing regime, between these limiting cases. Therefore, approaches that interpolate between these two limiting cases may allow a more accurate characterization of the magnetic inclusions at the source of non-exponential transverse relaxation in subcortical brain regions (Bauer and Nadler, 2002; Ziener et al., 2005). Moreover, brain tissue is inherently complex, involving a distribution of inclusions with varied sizes and susceptibilities originating from different cell types and structures. Some of these inclusions, such as larger cells like neurons, might be better described by the SDR, while others, such as smaller cells like glia, might be better described by the DNR. Additionally, the different models of the MRI signal considered here led to systematic differences in the estimates of the magnetic inclusions. Alternative models of the effect of the inclusions on the MRI signal should therefore be considered.

Future quantitative histological studies on cellular iron distributions in different subcortical areas may provide valuable priors for an informed choice of the appropriate model. In combination with the presented acquisition and fitting approach, this could enable the extraction of cellular characteristics non-invasively from non-exponential MR relaxometry.

### 5.3 Non-heme iron as a possible source of the non-exponential signal decay

Previous studies have suggested that the quadratic behaviour of non-heme iron may only be detectable at echo times well below 1 ms, below the range of achievable echo times, because clusters of non-heme iron were taken to be smaller than ∼100 nm (Yablonskiy et al., 2021). As a result, non-exponential signal decay was attributed to heme iron in deoxygenated blood. However, the results of the numerical simulations presented here (Figure 6) show that, in the SN, iron-rich dopaminergic neurons ∼15 *μ*m in size can lead to non-exponential decay in gradient-echo data acquired within a conventional range of echo times.

Additionally, while other magnetic materials like myelin or blood vessels also contribute to the non-exponential behaviour, the strongest deviations from the exponential decay occur in regions with high non-heme iron content (Section 5.1). This result further highlights the impact of cells rich in non-heme iron on the observed non-exponential decay behaviour.

## 6 Conclusions

In this study, we provided experimental evidence of non-exponential transverse relaxation signal decay in in vivo gradient-echo MRI data from subcortical brain regions at 3T. The behaviour of the decay is consistent with the effect of magnetic inclusions on the MRI signal predicted by theoretical studies. These theoretical models of the MRI signal yield improved fit with experimental data compared to the widely used exponential model. The larger deviations from exponential decay were observed in iron-rich subcortical regions (substantia nigra, globus pallidus). The experimental and numerical results presented here suggest that the observed non-exponential signal decay may originate from cells rich in non-heme iron such as dopaminergic neurons in the substantia nigra. From the estimates of the model parameters, we attempted to characterize the size, volume fraction and magnetic susceptibility of these cells. Non-exponential transverse relaxation signal decay provides new opportunities for the study of iron-related changes in neurodegenerative diseases non-invasively from MRI data, with increased specificity.

## 8 Author Contributions

RO: Conceptualization, Methodology, Investigation, Formal analysis, Writing-Original draft, Writing – Review and Editing, QR: Investigation, Writing – Review and Editing, EK: Investigation, Conceptualization, Writing – Review & Editing, VGK: Conceptualization, Writing – Review & Editing, IJ: Conceptualization, Writing – Review & Editing, AL: Conceptualization, Methodology, Investigation, Formal analysis, Writing-Original draft, Writing – Review and Editing, Supervision, Project administration, Funding acquisition.

## 10 Data and code availability

The code and data used for this analysis will be made publicly available upon publication.

## 12 Supplementary

**Figure S1.**
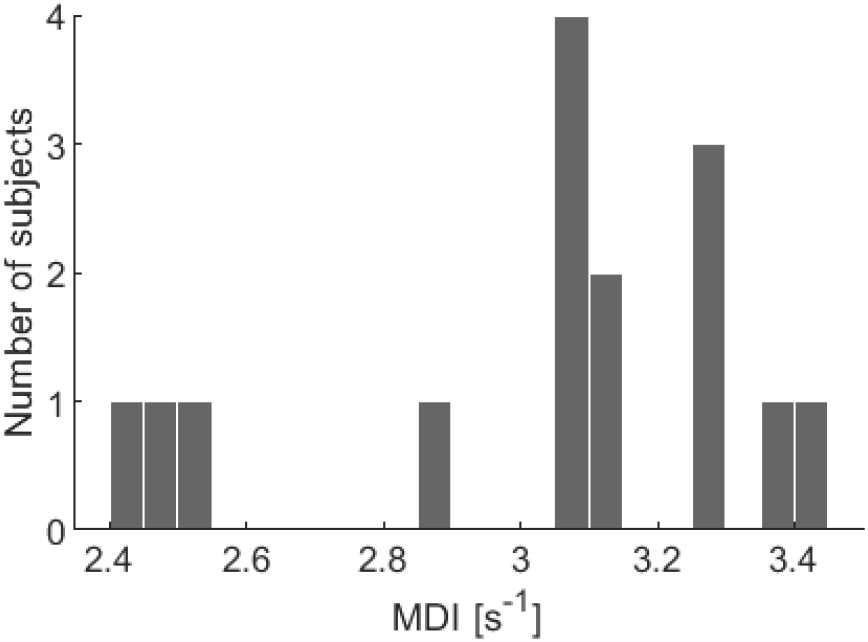
Distribution of Motion Degradation Index (MDI) across subjects and repetitions.

**Figure S2.**
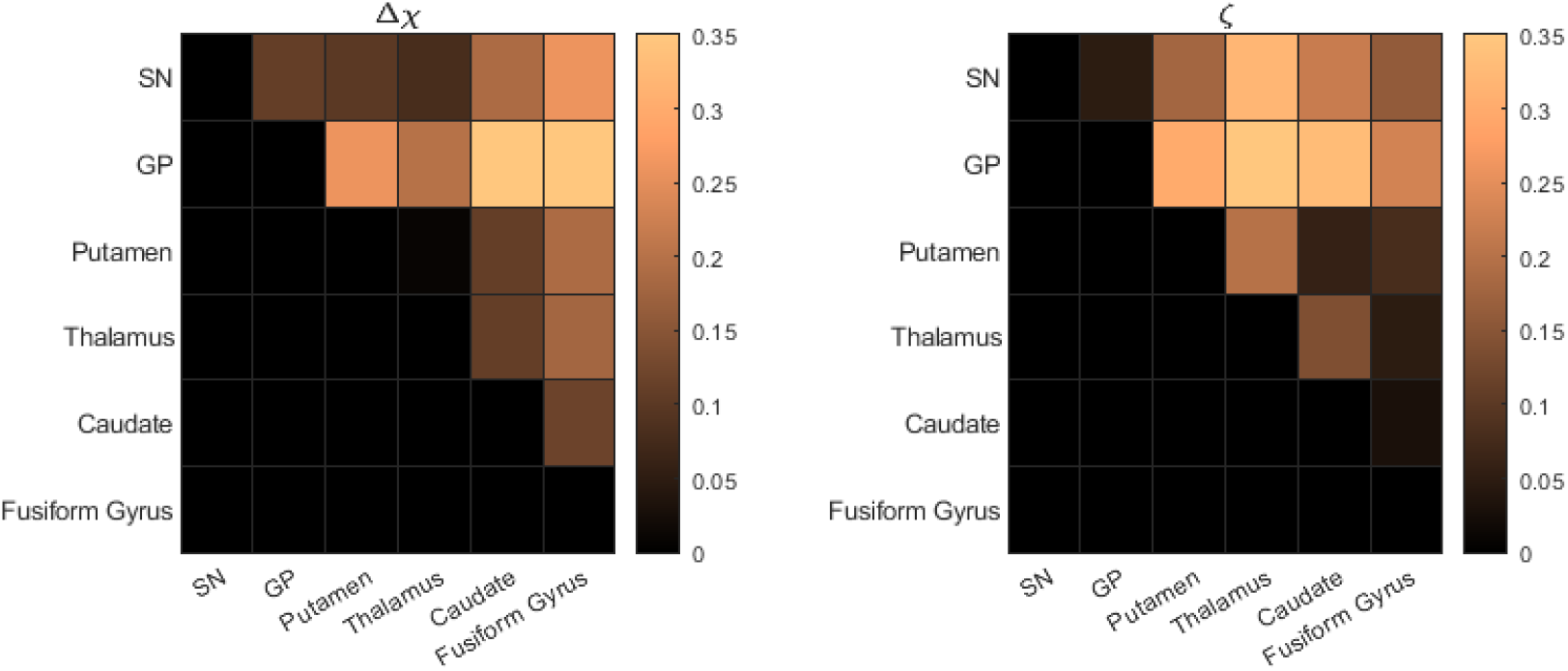
Absolute effect size (cliff’s delta) of the differences in Δ*χ* and *ζ* between subcortical regions for the AW models. A value of 0 suggests no difference between the two regions, while values closer to 1 indicate stronger associations. The p-values are not shown given that excluding the pair thalamus and putamen in Δ*χ*, all the remaining pairwise comparisons were significant (due to the large sample size). Note that given that the matrix is symmetric only the upper part is shown.

